# The Evolution of Bequeathal in Stable Habitats

**DOI:** 10.1101/336222

**Authors:** Parry M.R. Clarke, Mary Brooke Mcelreath, Brendan J. Barrett, Karen E. Mabry, Richard Mcelreath

## Abstract

Adults sometimes disperse while philopatric offspring inherit the natal site, a pattern known as *bequeathal*. Despite a decades-old empirical literature, little theoretical work has explored when natural selection may favor bequeathal. We present a simple mathematical model of the evolution of bequeathal in a stable environment, under both global and local dispersal. We find that natural selection favors bequeathal when adults are competitively advantaged over juveniles, baseline mortality is high, the environment is unsaturated, and when juveniles experience high dispersal mortality. However, frequently bequeathal may not evolve, because the fitness cost for the adult is too large relative to inclusive fitness benefits. Additionally, there are many situations for which bequeathal is an ESS, yet cannot invade the population. As bequeathal in real populations appears to be facultative, yet-to-be-modeled factors like timing of birth in the breeding season may strongly influence the patterns seen in natural populations.

## 1. INTRODUCTION

In this paper, we develop the first evolutionary models of *bequeathal*. Bequeathal is a type of breeding dispersal which occurs when a parent disperses to a new site, leaving a philopatric offspring to inherit the natal site and its resources. We know bequeathal occurs in nature from field studies of four mammal species: Columbian ground squirrels, *Urocitellus columbianus* (Harris and Murie 1984); kangaroo rats, *Dipodomys spectabilis* (Jones 1986); red squirrels, *Tamiasciurus hudsonicus* (Price and Boutin 1993, Berteaux and Boutin 2000); and woodrats *Neotoma macrotis* (Linsdale and Tevis 1951, Cunningham 2005). There is also evidence that it may occur in several other species: common wombats, *Vombatus ursinus* (Banks et al. 2002); hairy-nosed wombats (Johnson and Crossman 1991), *Lasiorhinus krefftii*; plateau pika (Zhang et al. 2017) *Ochotona curzoniae*; and wolverines (Aronsson and Persson 2018), *Gulo gulo*. Bequeathal has deep similarities with cooperative breeding and philopatric queuing (Kokko and Johnstone 1999, Kokko and Ekman 2002, Clutton-Brock 2006) in that related individuals cooperate to improve fitness outcomes, and juveniles stand to inherit the natal territory. The difference is that bequeathal does not involve group co-residence, and the costs of cooperation are paid by the dispersing adult rather than the offspring. Nonetheless, bequeathing adults often disperse short distances to nearby sites, where proximity to kin creates additional opportunities for cooperative behavior to evolve. In this light, bequeathal can be viewed as a type of cooperative breeding, and is part of the spectrum of strategies that help us understand the evolution of natal philopatry and kin cooperation (Clutton-Brock and Lukas 2012).

Despite bequeathal being empirically observed for nearly 70 years (Linsdale and Tevis 1951), there is no theoretical framework to explain its presence and absence. While natal dispersal is relatively well studied (Ronce 2007, Clobert et al. 2012), developing a greater understanding of bequeathal can teach us about the other side of the same behavioral coin, and adds a new dimension to our understanding of breeding dispersal (Paradis et al. 1998, Johst and Brandl 1999, Harts et al. 2016). Studying the exceptions to the norm in evolutionary ecology is often illuminating and can provide fresh insights into well-studied biological processes. As dispersal is a fundamental driver affecting the ecology, evolution, and population persistence of organisms (Bowler and Benton 2005), understanding the conditions which favor particular types of dispersal is of much importance.

Empirical studies of bequeathal are rare (Berteaux and Boutin 2000). However, reported rates of bequeathal are as high as 68% in red squirrels (Boon et al. 2008) and 30% in kangaroo rats (Jones 1986). All known examples of bequeathal occur in commodity-dependent species that require valuable resources such as a dens, burrows, middens, or resource caches for survival and reproduction (Lambin 1997). When resources critical to survival and reproduction are substantial and difficult to secure, parents may boost offspring fitness by bequeathing the natal site. Such offspring may stand a better chance of defending the natal territory than dispersing and acquiring a new one. Parents, on the other hand, are often in a better position to detect vacant territories or challenge existing ownership because of their enhanced experience and competitive skills. However, these benefits must be balanced against potentially high conflict between parents and offspring, especially in viscous populations (Kuijper and Johnstone 2012). This conflict of interest is inherent to a variety of social systems involving resource inheritance, such as cooperative breeding (Koenig et al. 1992), primatively eusocial and eusocial societies (Myles 1988).

Other factors, including parental age and condition, offspring size and competitive ability, territory quality, and population density, are thought to affect bequeathal (Price and Boutin 1993, Lambin 1997). However, with so many inputs, interpreting and synthesizing results from multiple studies is challenging. Some studies have found no relationship between parental age/condition and bequeathal (Price and Boutin 1993), while others have found an increase in bequeathal with age (Descamps et al. 2007). It seems clear that density matters, but how and when is unclear; bequeathal has been found to increase with local density in kangaroo rats (Jones 1986) but to decrease with density in Columbian ground squirrels and red squirrels (Harris and Murie 1984, Price and Boutin 1993, Boutin et al. 1993). Similarly inconsistent patterns of density-dependent dispersal have been observed across vertebrate taxa (Matthysen 2005).

Part of the problem in interpreting the current evidence is the lack of a general theoretical framework for understanding bequeathal dynamics (Berteaux and Boutin 2000). The problem of bequeathal lies at the intersection of parent-offspring conflict and dispersal, both long and large literatures (Trivers 1974, Godfray 1995, Hamilton and May 1977, Anderson 1989, Clobert et al. 2001, 2012). But very little work has directly addressed bequeathal. Price (1992) used dynamic programming to investigate optimal bequeathal for a single female, finding that timing of breeding was an important determinant of its adaptive value. But as the model did not include any population, just an individual female, it is difficultto interpret. Bequeathal, as a special form of dispersal, is inherently game theoretic, generating powerful frequency dependence. A game-theoretic model by Kokko and Lundberg (2001) comes closest to our target, in that it examines dispersal from and competition for territorial breeding sites, combined with conflict between an adult and a single offspring. However, their model examined residency in seasonal habitats with different productivity and survivorship, and it failed to find any bequeathal-like pattern among the evolutionarily stable strategies.

As a first step to building a theoretical framework for bequeathal, we present a simple bequeathal model. Our model considers parent-offspring conflict, competition for territories, local and global dispersal, and survival rates of adults and juveniles with overlapping generations. Like Kokko and Lundberg (2001), we consider production of a single offspring to avoid complications arising from sibling competition. This assumption is unrealistic in many cases, but allows for understanding of other factors before advancing to more complicated models. Unlike Kokko and Lundberg (2001) but like Hamilton and May (1977), we study a stable, uniform habitat, in order to eliminate many well-studied causes of dispersal in spatially and temporally variable environments. This is also unrealistic, but again allows for understanding the basic evolutionary logic of bequeathal, before studying it in stochastic environments, in which dispersal may be favored for other reasons.

A great deal of work remains to be done, extending these first models to consider facultative responses and additional strategies such as reproductive queuing. Still, even the simple models we analyze here are capable of producing a number of surprising dynamics. Therefore they are worth understanding in themselves before productive work can begin on extending them.

The major result of our analysis is that bequeathal is favored by the *comparative advantage* adults have in competing for sites. This advantage arises because there is more competition to acquire a new site than to retain an existing site. Since adults are better competitors, comparative advantage favors sending the better warrior to the most difficult battle. However, inclusive fitness considerations tend to work against bequeathal. Under clonal reproduction, the adult and juvenile will agree that the best warrior serve in the harshest battle. But since adults and juveniles are imperfectly related, they disagree, under some range of costs and benefits. Any factor that reduces the adult’s costs will therefore help bequeathal evolve. Such factors include adults having high mortality risk and low residual reproductive value, such as at the end of life. Conversely, any factor that reduces juvenile benefits will work against bequeathal. For example, if juveniles are fragile, having high baseline mortality, then it makes little sense to bequeath territory to them. We outline the mathematical argument that leads to these conclusions, ending the paper with a discussion of un-modeled factors that may also strongly influence the facultative use of bequeathal in natural populations.

## 2. Model definition

We use a mix of methods—including formal analysis, numerical sensitivity analysis, and individual-based simulation—to construct and understand our models of bequeathal. We begin by defining the global and local dispersal models analytically. Table 1 summarizes the symbols used in the models, each of which is explained in the following sections.

**Table 1.**
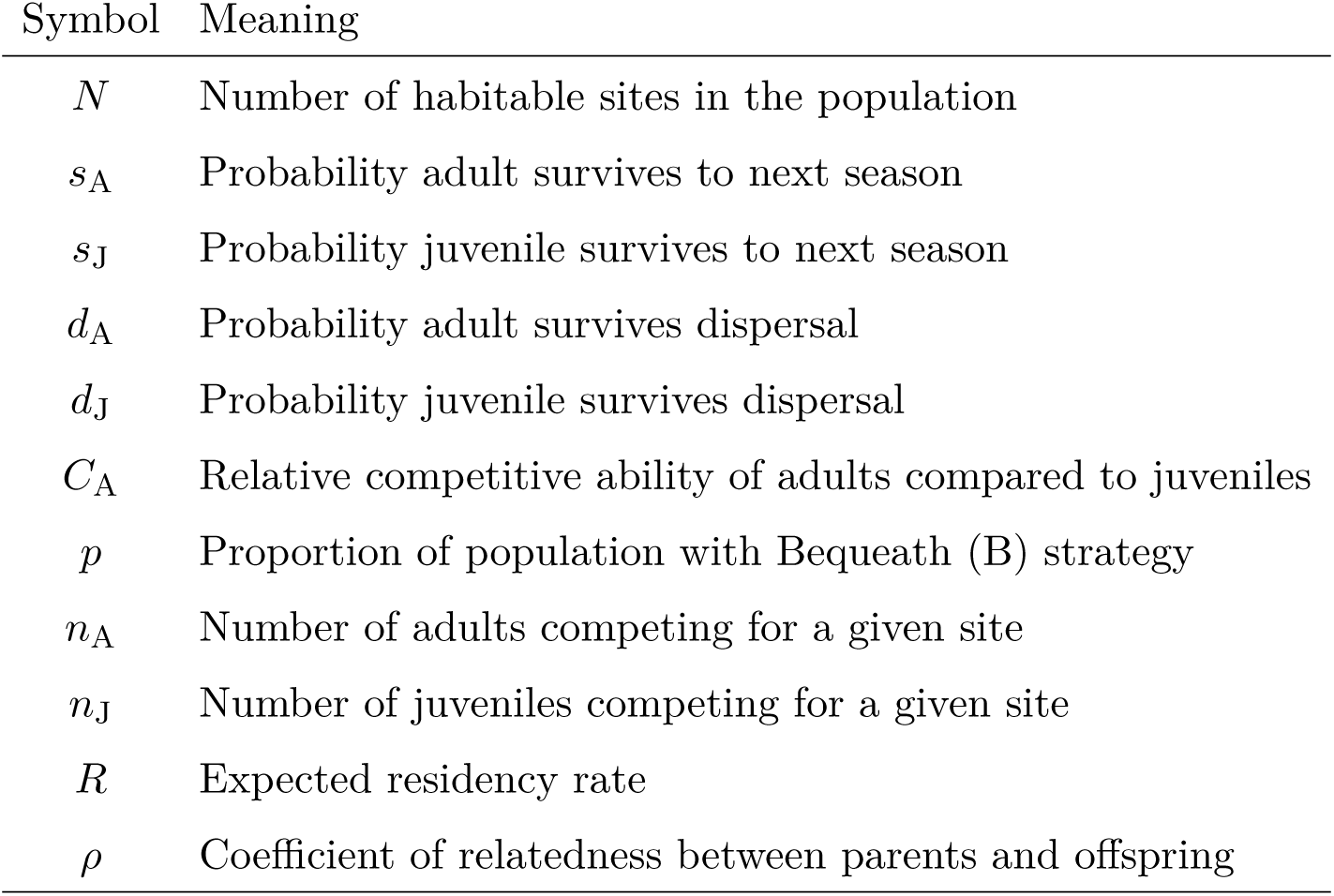
Symbols used, with their meanings.

### 2.1 Population and life cycle

Imagine a population of organisms with overlapping generations, living at *N* spatially separated sites. Only one adult can survive and reproduce at each site, and each adult produces one same-sex (female) juvenile offspring each breeding season. The life cycle proceeds in the following sequence: 1) birth of offspring, 2) dispersal of either the offspring or parent, 3) competition for site occupancy, and 4) probability of survival to the next breeding season.

Juveniles reproductively mature in one breeding season. At the end of each breeding season, adults and juveniles may die, prior to reproduction in the next season. Let *s*_A_be the probability a resident adult (A) survives to the next breeding season. Let *s*_J_ be the corresponding juvenile survival probability. When *s*_A_ = *s*_J_ = 1, all sites will remain occupied. The environment will be saturated. When either survival probability is less than one, some open sites may exist. Thus these models allow us to examine the effects of saturation and open environments, as emergent properties of vital parameters, rather than exogenous assumptions.

### 2.2 Heritable strategies

Assume reproduction is sexual and haploid. Also assume two pure heritable strategies, Bequeath (B) and Stay (S). Both strategies are expressed in adults. A bequeathing adult always disperses after reproduction, arriving at an “away” site. This leaves its offspring behind to compete to retain the natal “home” site. A staying adult always evicts its offspring, forcing it to compete for an “away” site, while the adult remains behind to compete to retain the “home” site.

We have also analyzed an infinite-allele model that allows continuously varying strategies between pure Bequeath and pure Stay, using a heritable probability of bequeathing. The continuous strategy space produces the same results, in this case, owing to a lack of geometric mean fitness effects (bet hedging), stable internal equilibria, and evolutionary branching. Therefore we stick to the discrete strategy case in this paper, for ease of understanding. The individual-based simulation code we include in the Supplemental can be toggled to continuous strategy space for comparison.

### 2.3 Dispersal

We have analyzed two extreme dispersal models, a global model and a local model. In the global model all sites are equidistant; consequently, dispersal from any site has an equal probability of arriving at any other site. In the local dispersal model, sites are arranged in a ring, and individuals can disperse only to one of two neighboring sites, at random. Real dispersal patterns are probably intermediate between these two extremes.

We assume that dispersal is costly, carrying a chance of dispersal-related mortality. These costs may be due to increased predation risk during dispersal, energetic costs, or limited knowledge of resource availability in new sites. Let *d*_A_ be the probability that an adult survives dispersal and arrives at a new site. Let *d*_J_ be the probability that a juvenile survives dispersal. Typically, *d*_A_ *> d*_J_, and so we focus on that condition, considering whether it is necessary or not for Bequeathal to be an ESS.

### 2.4 Competition

Individuals must compete to retain or colonize sites. All individuals who disperse into or remain in a site compete for it. We assume a lottery-type competitive model, in which all individuals arriving or residing at a site simultaneously compete for it. Adults have an advantage over juveniles in competition, and we express this advantage as a relative advantage *C*_A_ *>* 1. The probability that an adult retains or occupies a site with *n*_A_ other adult competitors and *n*_J_ juvenile competitors is:

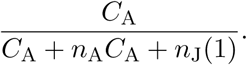

After competition, a single individual survives to occupy each site.

### 2.5 Expected fitness

Using the assumptions above, we can write expected inclusive fitness expressions for B and S. We fully develop the global dispersal model first, before specifying how the local model differs. The global model can be derived for any population frequency of Bequeath, *p*, while the local model cannot. However, both models can be analyzed for the ESS conditions of both B and S.

#### 2.5.1 Global dispersal and fitness

Let *p* be the proportion of the population with strategy B. Let *R* be the proportion of sites with a resident adult, at the start of each breeding season. The goal is to compute the probability *n*_A_ adults and *n*_J_ juveniles immigrate to a particular site. Under the assumption that dispersal events are independent of one another, the probability that *n*_A_ adults and *n*_J_ juveniles arrive at a particular site will be multinomial with three categories (adult, juvenile, none) and *N -* 1 trials. As the number of sites *N* grows large, the distribution approaches a bivariate Poisson, just like a binomial distribution with low probability approaches univariate Poisson as the number of trials becomes large. Therefore in the limit *N → ∞*:

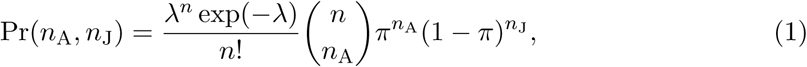

where λ = *R* (*pd*_*A*_ +(1−*p*)*d*_*J*_)is the average number of immigrants (either adult or juvenile) entering the site, *n* = *n*_A_ + *n*_J_, and

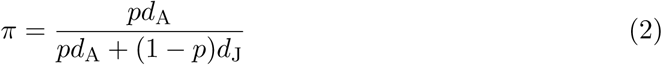

is the proportion of the surviving dispersal pool that is adult. Pr(*n*_A_, *n*_J_) is just a special case of a multivariate Poisson process, with uncorrelated dimensions. But it can be motivated more easily by considering that dispersal events are independent Poisson samples that are equally likely to arrive at the focal site. Whether a disperser is adult or juvenile can then be viewed as a binomial process, independent of arrival. Note that were adults and juveniles to use different dispersal strategies, varying in distance or some other aspect, then some other function would be required.

The expected residency rate *R* is dynamic, but quickly reaches a steady state expectation. The steady state of *R* is defined implicitly by the recurrence:

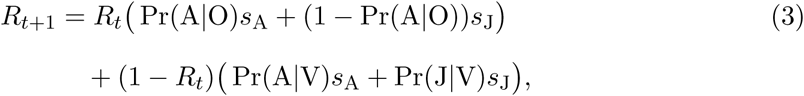

where *R*_*t*_ is the proportion of sites that are occupied at time *t* and Pr(A*|*O) is the probability an adult (A) wins a site that is occupied (O). Similarly, Pr(A*|*V) is the probability an adult wins a vacant (V) site. Pr(J*|*V) is the probability a juvenile (J) wins a vacant site. This recurrence cannot in general be solved explicitly for the steady-state value of *R*, the value that makes *R*_*t*+1_ = *R*_*t*_. But it can be solved numerically. A Mathematica notebook (Wolfram Research Inc. 2010) that computes *R*, as well as all of the other numerical results to follow, can be found through a link in the Supplemental Materials.

Using the above definitions, we can write the expected fitness of the Bequeath (B) and Stay (S) alleles. There are two components to this fitness measure. The first is the probability of retaining the home (natal) site. For a Bequeath individual, this is:

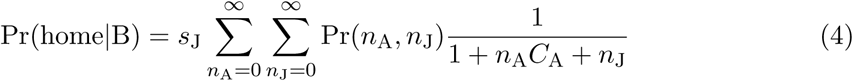

The juvenile stays at the site, competing with *n*_A_ adult immigrants and *n*_J_ juvenile immigrants. The juvenile survives the season with probability *s*_J_.

The other component of fitness is the probability of acquiring the away site to which the adult disperses. This is:

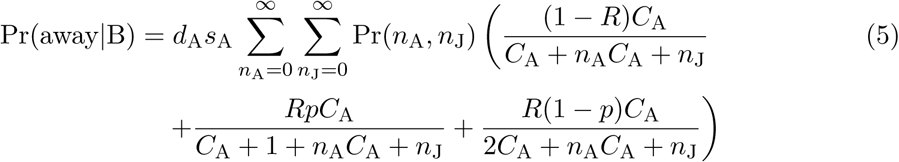

If the bequeathing adult survives dispersal, it competes with a resident *R* of the time, in addition to another *n*_A_ adult immigrants and *n*_J_ juvenile immigrants. Since the number of sites is very large, the distribution of immigrants here is the same as before, not conditional on the focal immigrant, because dispersal events are independent in the Poisson process. If the number of sites were small, or dispersal were local, this would not be true, as we explain later.Finally, we devalue fitness from the offspring, due to imperfect inheritance. This gives us inclusive fitness:

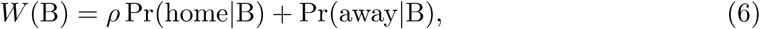

where *ρ* is the coefficient of relatedness between the adult and juvenile. For a typical example, this would be *ρ* = 0.5. But for a maternally inherited trait, it might be *ρ* = 1.

The fitness expression for the Stay strategy is constructed similarly:

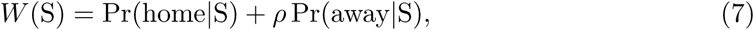

where:

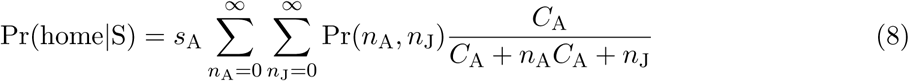

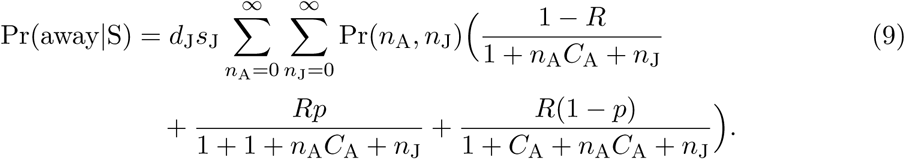

Note that while the expressions *W* (B) and *W* (S) are presented as inclusive fitness expressions, they are just expected growth rates. No weak selection approximation or other assumptions typical of other inclusive fitness models have been made.

#### 2.5.2 Local dispersal and fitness

The local dispersal model is analogous. However, the probability Pr(*n*_A_, *n*_J_) under local dispersal cannot be approximated by a Poisson distribution, even at *N → ∞*, because at most two sites (neighbors) contribute dispersers to any focal site. Additionally, the disperser pool is no longer independent of a focal disperser arriving at an away site. Furthermore, it is not easy to specify the distribution of immigrants for any population frequency of Bequeath, *p*, because local dispersal generates spatial correlations in genotypes—the population residency rate *R* will not tell us the relevant residency probability at every locale.

It is possible, however, to completely define the model for invading B and invading S, that is for 𝒫 *≈* 0 and 𝒫 *≈* 1. This allows us to conduct standard ESS analysis, even though we will not be able to find the location of any internal equilibria. This turns out to be sufficient for this model. But we have also verified all of these inferences using individual-based simulation, which is available through the link in Supplemental Materials.

Constraining *p ∈ {*0, 1*}*, the distribution of immigrants is now defined by a simple binomial process, as each neighboring site contributes an immigrant half of the time (it can go in either direction), discounted by the probabilities of residency *R* and dispersal survival *d*_A_ and *d*_J_. In other words, each immigrant is a coin flip from a biased coin with probability of arrival of *π* = *R*(*pd*_A_ + (1 − *p*)*d*_J_).

Whether there are one or two “coins” to flip depends upon our focus. When focusing on a home site, there are two neighbors who may contribute immigrants. But when focusing on an away site, the focal disperser counts as one of the neighbors, and so there is only one “coin” to flip. With these facts in mind, we can define inclusive fitness much as before. The expressions add little insight in themselves, and so we include them only in the appendix. The Mathematica notebook in the Supplemental contains all of these expressions and computes fitness differences from them.

## 3 Model Results

There are two antagonistic forces that strongly influence when Bequeath can be an ESS. The first is the *comparative advantage* that adults have in competition. This advantage favors Bequeath. The second force, opposed to the first, is the *conflict of interest* between parent and offspring that arises from sexual reproduction. Baseline survival, dispersal survival, and dispersal pattern (local or global) all interact with these two forces.

Even a model as simple as this one is very complex. Therefore we explain these two antagonistic forces first, without reference to dispersal pattern or baseline and dispersal survival rates. We consider how local and global dispersal differ, through their effects on comparative advantage and conflict of interest. Then we vary adult and juvenile survival rates to show how they interact with adult comparative advantage and parent-offspring conflict of interest.

### 3.1 Bequeathal is favored by comparative advantage

Assume for the moment that *s*_A_ = *s*_J_ = 1 and that *d*_A_ = *d*_J_ = 1 so that there is no baseline nor dispersal mortality. As can be seen by substituting these values in Equation 3, these assumptions imply that all sites are always occupied (*R* = 1), a saturated environment. Figure 1 illustrates the nature of invasion and stability under these conditions. Each of the four diagrams in Figure 1 illustrates movement from and into a focal “home” site for a rare invader, as well as movement from and to an “away” site the invader attempts to claim. This is a cartoonish representation of the full model, but will serve to explain the basic forces in the model, before moving on to nuances.

**FIGURE 1.**
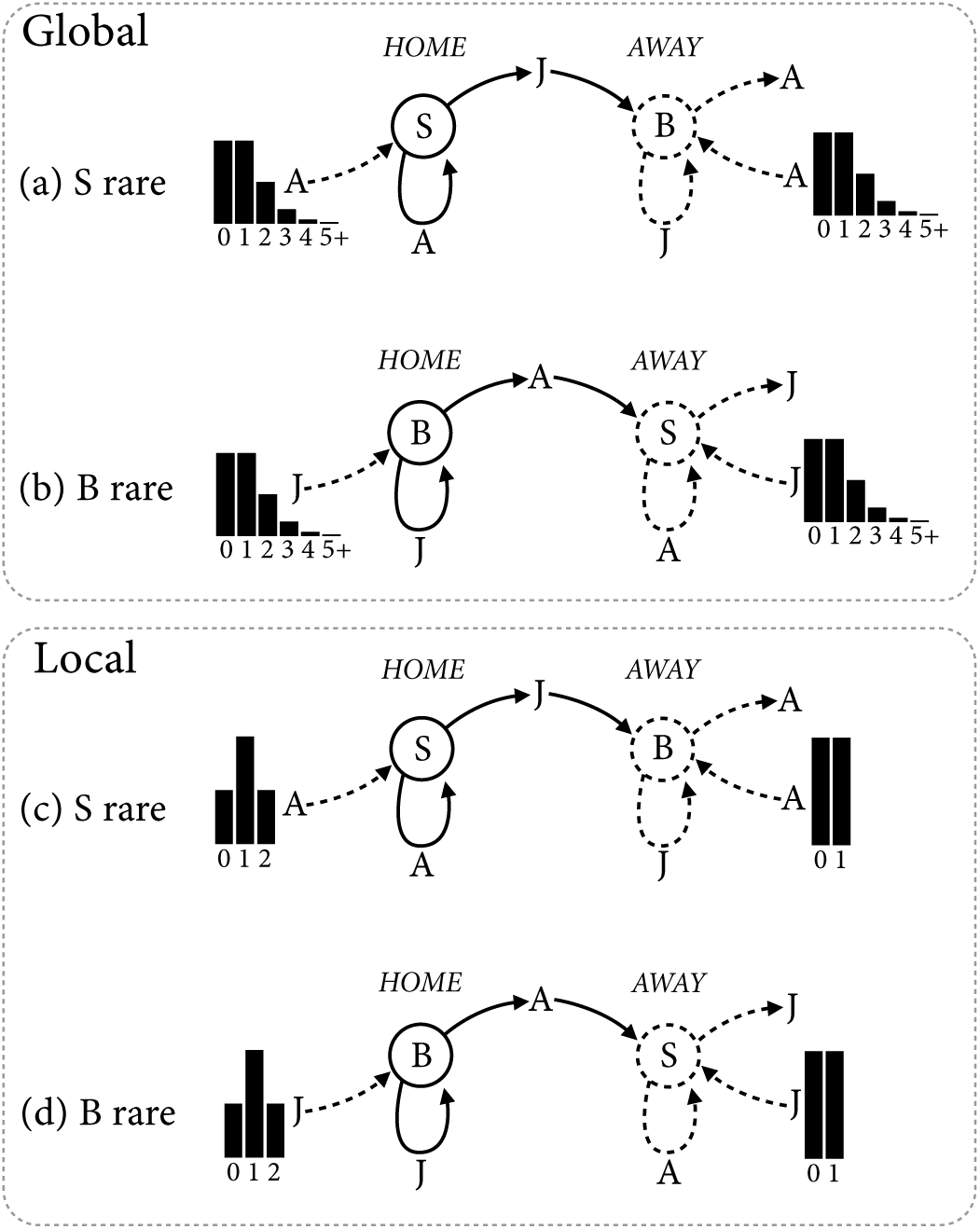
Schematic of invasion and stability in the global and local dispersal models, for *s*_A_ = *s*_J_ = 1 and *d*_A_ = *d*_J_ = 1. B indicates Bequeath and S indicates Stay. Solid lines indicate invader (rare type) and dashed lines resident (common type). The clusters of black bars in each sub-figure represent probability distributions of immigrants, with numbers of immigrants labeled along the bottom. Top two schematics: Bequeath stability (top) and invasion (bottom), under global dispersal. Bottom two schematics: Local dispersal.

Consider only the top two diagrams for now, (a) and (b). These diagrams represent Bequeath’s stability (a) and invasion (b) under the global dispersal model. When B is common (top), a lone S adult (A) remains at its home site, while evicting a juvenile (J) to disperse to an away site. The home site receives dispersers from other sites, all of which are occupied by B, and so all of the immigrants to the home site are adults (A). At the away site, the juvenile S individual is joined by adults dispersing from other sites to compete with a resident juvenile B individual. The distributions of immigrants at both the home and away sites are the same Poisson distribution, with a mean of 1, because of the global dispersal pattern.

Now consider the amount of competition at home and away. To retain the home site, the adult S individual competes with, on average, 1 other adult. Any competitive advantage of adults has no effect here, because all immigrants are adult, when B is common. In contrast, to acquire the away site, the lone juvenile disperser competes with a juvenile resident and, on average, 1 adult immigrant. Therefore there is one additional competitor at the away site, and the juvenile must contend with its disadvantage against an adult (assuming *C*_A_ *>* 1). So, Stay sends its juvenile to an away site at which it must compete against, on average, one additional juvenile. Also, any competitive advantage of adults hurts Stay, because as *C*_A_ increases, the chance of acquiring the away site decreases. For very large *C*_A_, the only way for a S juvenile to acquire an away site is for no adults to immigrate.

The situation is nearly reversed when Bequeath invades, as shown in Figure 1(b). Now a B juvenile remains home and competes with, on average, 1 other juvenile. Immigrants are all juvenile now, because Stay is common. Competitive advantage of adults (*C*_A_ *>* 1) is again irrelevant for the invader retaining the home site. But at the away site, the dispersing B adult does better as *C*_A_ increases, since its competitive advantage reduces the impact of any immigrant juveniles. If *C*_A_ = 1 the dispersing adult acquires the away site one-third of the time, on average. But for very large *C*_A_, it will acquire the away site one-half of the time.

Considering both Figure 1(a) and Figure 1(b) together, the principle reason that Bequeath can be an adaptation is that it uses the comparative advantage of adults by allocating the better warrior, the adult, to the worse battlefield, the away site. In contrast, Stay allocates the worse warrior, the juvenile, to the worse battlefield. In the mathematical appendix (Equations A1 - A5a,b), we show that, as long as no other forces are in play (*C*_A_ *>* 1 and *ρ* = 1), Bequeath is always an ESS and Stay is never an ESS.

The same principle applies to the local dispersal model, illustrated by Figure 1(c) and (d). However, the excess competition at away sites, compared to the home site, is smaller than in the global dispersal model. This fact has no impact on the long run dynamics, as long as *ρ* = 1. B is still favored by comparative advantage and uniquely an ESS. So we postpone discussion of local dispersal until the next section.

### 3.2 Sexual reproduction and conflict of interest

The principle of comparative advantage will not uniquely determine the evolutionary result, unless the juvenile and adult have no conflict of interest. When *ρ* = 1, there is no conflict of interest, and selection favors allocating the adult to the more dangerous away site. The adult and juvenile always agree. But for *ρ <* 1, there is a conflict of interest, with bequeathal representing a costly action by the adult. As *ρ* gets smaller, selection favors adults choosing the easier battle, which is always the home site. However, large *C*_A_ can compensate, allowing B to continue to be stable, even when *ρ* is so small that B can no longer invade the population.

To appreciate how conflict of interest and comparative advantage interact, in Figure 2 we map regions of stability for B and S for combinations of *ρ* and *C*_A_. Focus for now on only the upper-left, panel (a), the enlarged plot with labeled regions. The horizontal axis is the magnitude of *C*_A_, expressed as the base-2 logarithm, a “fold” value. If you folded a piece of paper in half 10 times, then its thickness would be 2^10^ layers, a 10-fold increase in thickness. Likewise you can read the value log_2_ *C*_A_ = 10 as a 10-fold increase in adult competitive ability, relative to a juvenile. The vertical axis is *ρ*, from complete conflict at the bottom to complete agreement at the top. The colored regions represent different combinations of *ρ* and *C*_A_ for which B and S are not evolutionarily stable. In the orange regions, S is not an ESS. In the blue regions, B is not an ESS. In the white region, both B and S are evolutionarily stable. The red and blue curves show the boundaries for the different dispersal models, with global dispersal represented by the solid curves and local by the dashed.

**FIGURE 2.**
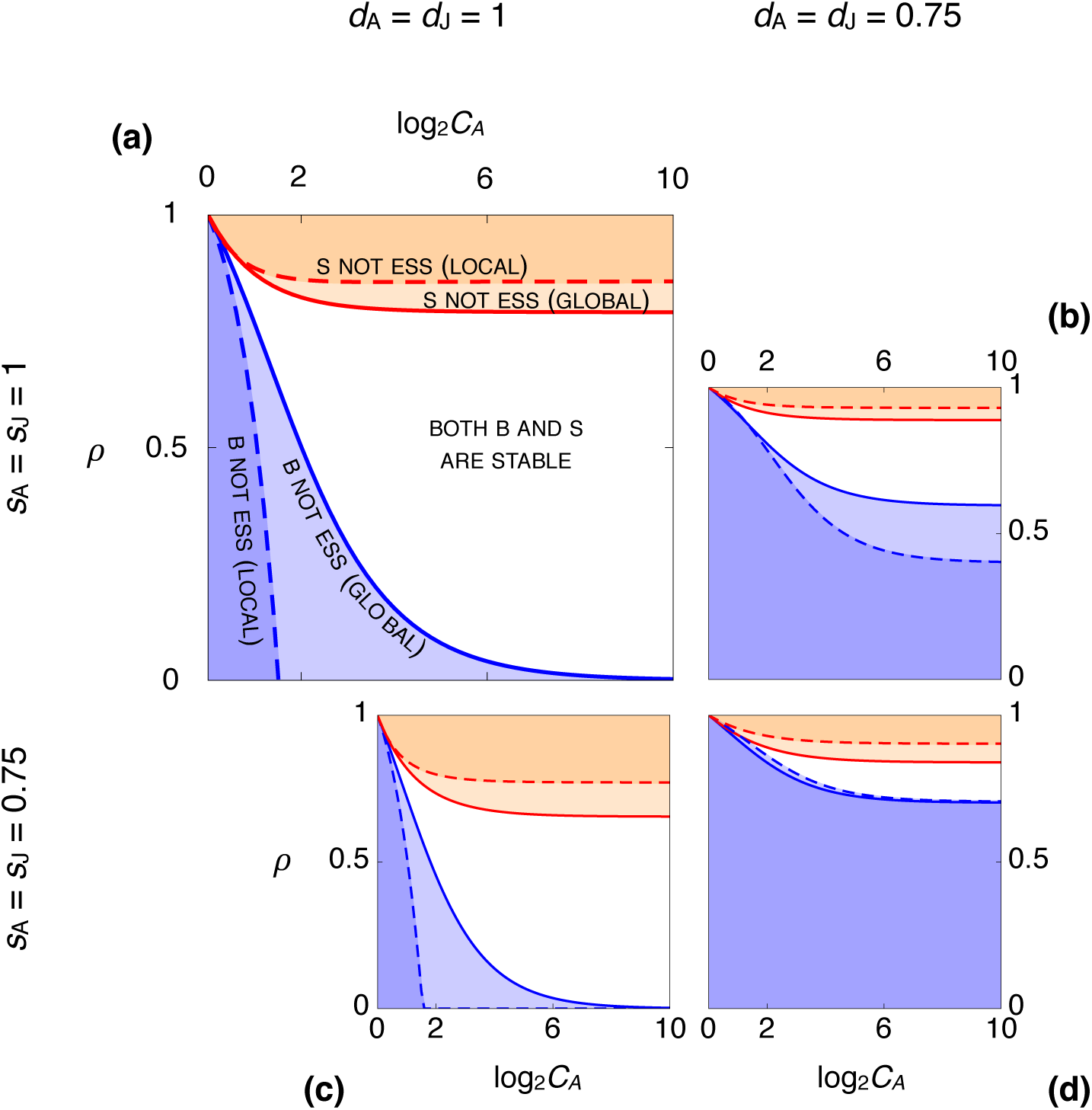
Effects of relatedness, *ρ*, and adult competitive ability, *C*_A_, on stability of Bequeath and Stay. In each panel, horizontal axis is the logarithm of *C*_A_ and vertical axis is *ρ*. The shaded regions indicate combinations of *C*_A_ and *ρ* for which either Bequeath (orange) or Stay (blue) is the only ESS. In the white region, both Bequeath and Stay are ESSs. Boundaries for the global dispersal model are solid. Boundaries for the local dispersal model are dashed. Each panel shows regions for different combinations of dispersal survival and baseline survival. Top row: *s*_A_ = *s*_J_ = 1. Bottom row: *s*_A_ = *s*_J_ = 0.75. Left column: *d*_A_ = *d*_J_ = 1. Right column: *d*_A_ = *d*_J_ = 0.75.

In Figure 2(a), there is no dispersal mortality nor baseline mortality. At the top, the results correspond to the inferences in the previous section: the comparative advantage of adults renders Bequeath an ESS (and Stay not an ESS) for all *C*_A_ *>* 1 (log_2_ *C*_A_ *>* 0). But as *ρ* decreases, the orange regions become increasingly restricted to large *C*_A_ values. By the time *ρ* reaches 0.5, corresponding to sexual reproduction, either only S is an ESS (blue regions) or both B and S are ESSs. At the limit *ρ* = 0, B is never an ESS, although if *C*_A_ is large enough, even tiny amounts of relatedness are sufficient for B to be an ESS. We prove this result in the appendix.

To understand these results, consider Stay to be a “selfish” strategy while Bequeath is “cooperative.” A Bequeath adult disperses at a personal cost, because there is more competition at the away site, leaving the easier home site for the juvenile to defend. When *ρ* = 1, the interests of the adult and juvenile are completely aligned, and so the adult favors the strategy that results in the greatest joint success (family growth). But when *ρ <* 1, the adult and juvenile will disagree.

Provided *C*_A_ is large enough, B can remain stable. But for small *C*_A_, B may not be an ESS. The reason B can be stable even when it cannot invade is because of positive frequency dependence. When B is rare, the adult is dispersing into a site with a resident S adult, in addition to any juvenile immigrants from other sites. For an adult, competing against another adult for the away site is much harder than defending the home site from invading juveniles. But as B increases in frequency, more and more away sites are occupied by juveniles left behind by B adults. It is simultaneously true that more adults enter the dispersal pool, and so adults invade the away site. But this effect happens at both the home and away site and so does not affect the relative cost of adult dispersal. This means that Bequeath does better the more common it becomes, because the away site becomes easier to win, reducing the costliness of adult dispersal.

The boundaries for global and local dispersal, shown by the solid and dashed curves, sometimes differ greatly. The major effect of local dispersal is to make it harder for either strategy to invade the population. Local dispersal makes the white region larger, and so more combinations of parameters lead to both B and S being evolutionarily stable. To understand why, it is helpful to refer again to Figure 1. Under local dispersal, at most 2 individuals can immigrate into any site. Therefore, while the average number of immigrants remains the same as in the global model, the distribution is different. First, the probability of zero immigrants at the home site is reduced under local dispersal. Under global dispersal, the probability of zero immigrants is exp(*-*1) *≈* 0.37, while under local dispersal it is only 0.25 (the chance of two coin flips coming up tails). This makes the effective amount of competition greater under local dispersal. Second, the focal disperser now counts for one of the immigrants at the away site. So a rare strategy disperser now competes against, on average, one resident and one-half immigrant, instead of one resident and one immigrant, as under global dispersal. Indeed, the probability of no additional immigrants at the away site has increased to 0.5 under local dispersal, in contrast to 0.37 under global dispersal.

This reduced competition at the away site and increased competition at the home site helps Bequeath, by reducing the effective cost of adult dispersal. It is still true that average competition at the away site is greater than average competition at home. But a smaller difference under local dispersal means that B can be stable for smaller values of *ρ* than it can under global dispersal. Simultaneously, Stay becomes stable under local dispersal for larger values of *ρ*. The sword of local dispersal cuts both ways: a smaller cost for a dispersing adult is also a smaller benefit for a resident juvenile. This means that Bequeath gains less under local dispersal than it does under global, resulting in both dashed boundaries in Figure 2(a) receding and increasing the range of conditions for which both B and S are ESSs.

The other plots in Figure 2 show the interaction of *ρ* and *C*_A_ under different values of dispersal and baseline survival. In (c), baseline survival for both adults, *s*_A_, and juveniles, *s*_J_, is reduced by 25%. This creates open habitat, effectively reducing competition at the away site. Under global dispersal, the steady state residency becomes *R ≈* 0.56. Under local dispersal, *R ≈* 0.61. Competition at the home site is also reduced, as fewer other sites have residents to produce immigrants. But this reduction in the disperser pool applies equally to home and away sites. In aggregate, lowered baseline survival benefits Bequeath, by reducing the relative intensity of competition at the away site. This results in an increased orange region, a reduction of the region in which Stay can be an ESS.

In Figure 2(b), we instead reduce dispersal survival by 25%, setting *d*_A_ = *d*_J_ = 0.75. Dispersal mortality has the opposite effect, to aid Stay over Bequeath. Unlike a reduction in baseline survival, a reduction in dispersal survival does not necessarily result in open habitat. Here, the environment remains saturated at *R* = 1. Since residents always survive, as long as any individual arrives at a site, the site will remain occupied, eventually filling the environment. Now the cost of dispersal is greatly increased. If *ρ* = 1, this has no effect, because the adult will still agree to disperse, since both the adult and juvenile must pay the same dispersal cost (25%). But as long as *ρ <* 1, the cost quickly becomes too great for the adult, favoring Stay. The region in which B can be an ESS is greatly reduced.

Combining 25% baseline and dispersal mortality, in panel (d), demonstrates a strong interaction between these two forms of mortality. To further understand the effects of the mortality parameters, we proceed in the next sections by fixing *ρ* = 0.5, representing sexual reproduction, and allowing adult and juvenile survival rates to vary independently.

### 3.3 Baseline mortality

Figure 3 shows the effects of independently varying adult and juvenile baseline mortality, for zero dispersal mortality (top row) and 20% dispersal mortality (bottom row), for two values of *C*_A_ (left and right columns). Relatedness is set to *ρ* = 0.5. Colors have the same meanings as before. The purple region in the lower-left of (c) and (d) indicates combinations of parameters at which a population of Stay individuals is non-viable, approaching a residency rate *R* = 0. The conditions for viability are *s*_A_ + *p*_J_*s*_J_ *>* 1, for a monomorphic population of Stay, and *s*_J_ + *p*_A_*s*_A_ *>* 1 for a monomorphic population of Bequeath. Note that Stay can be both an ESS and non-viable, as sometimes happens in models with both ecological and evolutionary dynamics. Also note that the conditions for viability refer to expectations. Many parameter combinations will lead to extirpation with high probability, even when they strictly satisfy the conditions above. Populations near the purple region are highly endangered.

**FIGURE 3.**
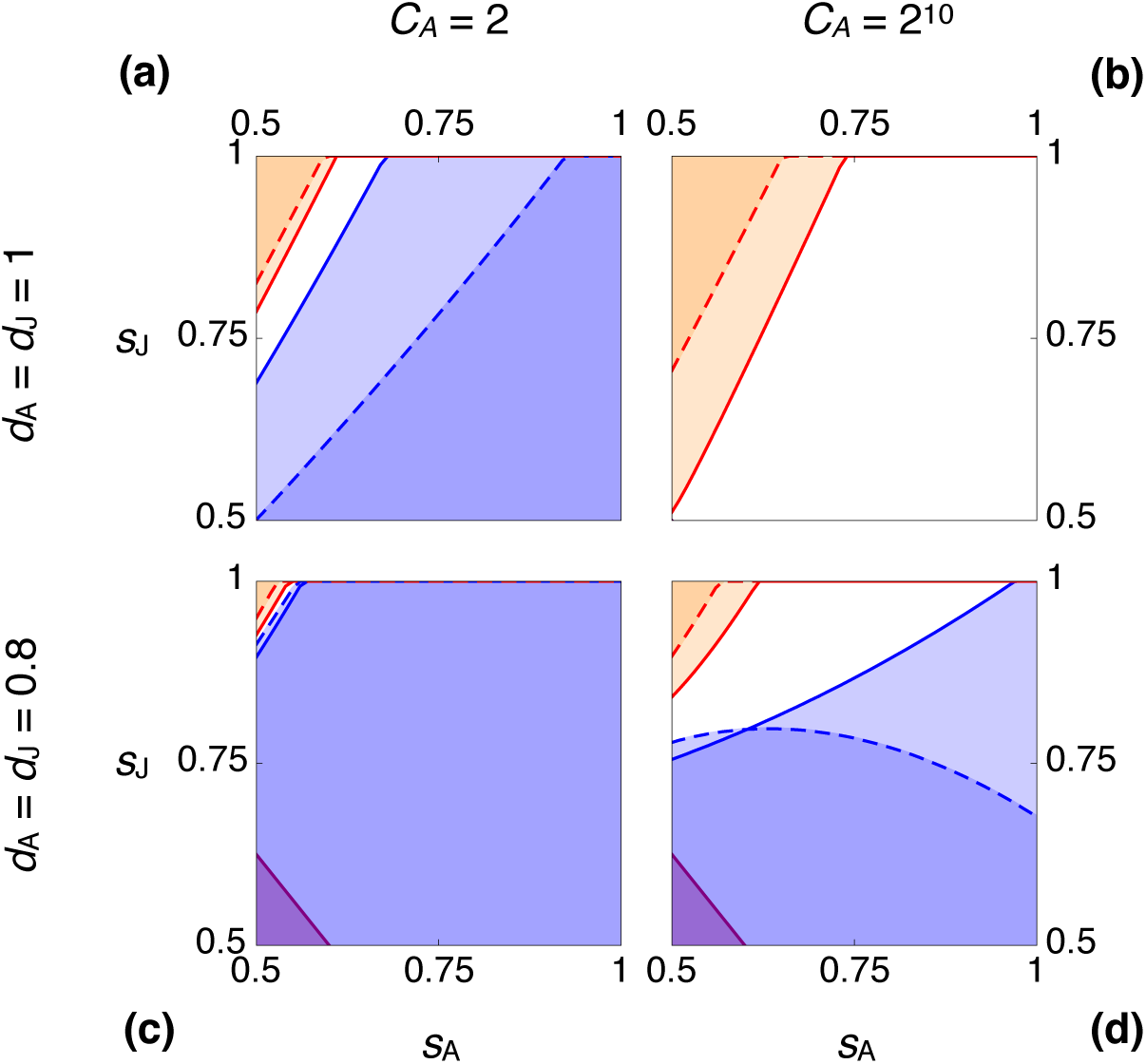
Effects of adult and juvenile baseline survival, *s*_A_ and *s*_J_, on stability of Bequeath and Stay. Colors and curves as in Figure 2, but now with *ρ* = 0.5 only. The purple region in the lower-left of (c) and (d) indicates parameter combinations leading to extirpation. In each panel, horizontal axis is adult survival and vertical axis is juvenile survival. Top row: *d*_A_ = *d*_J_ = 1. Bottom row: *d*_A_ = *d*_J_ = 0.8. Left column: *C*_A_ = 2. Right column: *C*_A_ = 2^10^.

Perhaps counterintuitively, Bequeath does best when adult survival, *s*_A_, is low while juvenile survival, *s*_J_, is high. When adult survival is low, the residual reproductive value of an adult is also low. This effectively reduces the cost to the adult of bequeathing the home site. Since the adult will likely die anyway, better for it to provide a benefit to the offspring. However, unless *s*_J_ is also sufficiently large, the juvenile will not live to enjoy any bequeathed benefit. As a result, the orange regions lie in the upper-left corner of each plot in Figure 3.

Adding dispersal mortality (bottom row) and increasing adult competitive advantage (right column) have the same effects as before. Dispersal mortality reduces the region in which Bequeath can be stable. But large *C*_A_ can compensate, increasing both the region in which B is the only ESS (orange) and especially the region in which both B and S are ESSs (white). The effect of increasing *C*_A_ on the stability of Bequeath is pronounced for the local dispersal model, as seen in Figure 3(d), where B is stable for most of the total plot, but only when dispersal is local (the dashed boundaries).

### 3.4 Dispersal mortality

Figure 4 shows the effects of independently varying adult and juvenile dispersal mortality, for zero baseline mortality (top row) and 20% baseline mortality (bottom row), for two values of *C*_A_ (left and right columns). Again, *ρ* = 0.5 in all four plots. Intuitively, Bequeath does best when adult dispersal survival, *d*_A_, is large and juvenile dispersal survival, *d*_J_, is low. Such an asymmetry further improves the adult’s comparative advantage, by increasing the relative probability that the adult will reach the away site. However, as long as *C*_A_ is large enough, there are many combinations *d*_A_ *< d*_J_ at which Bequeath is stable, even though it cannot invade (white regions in the figure).

**FIGURE 4.**
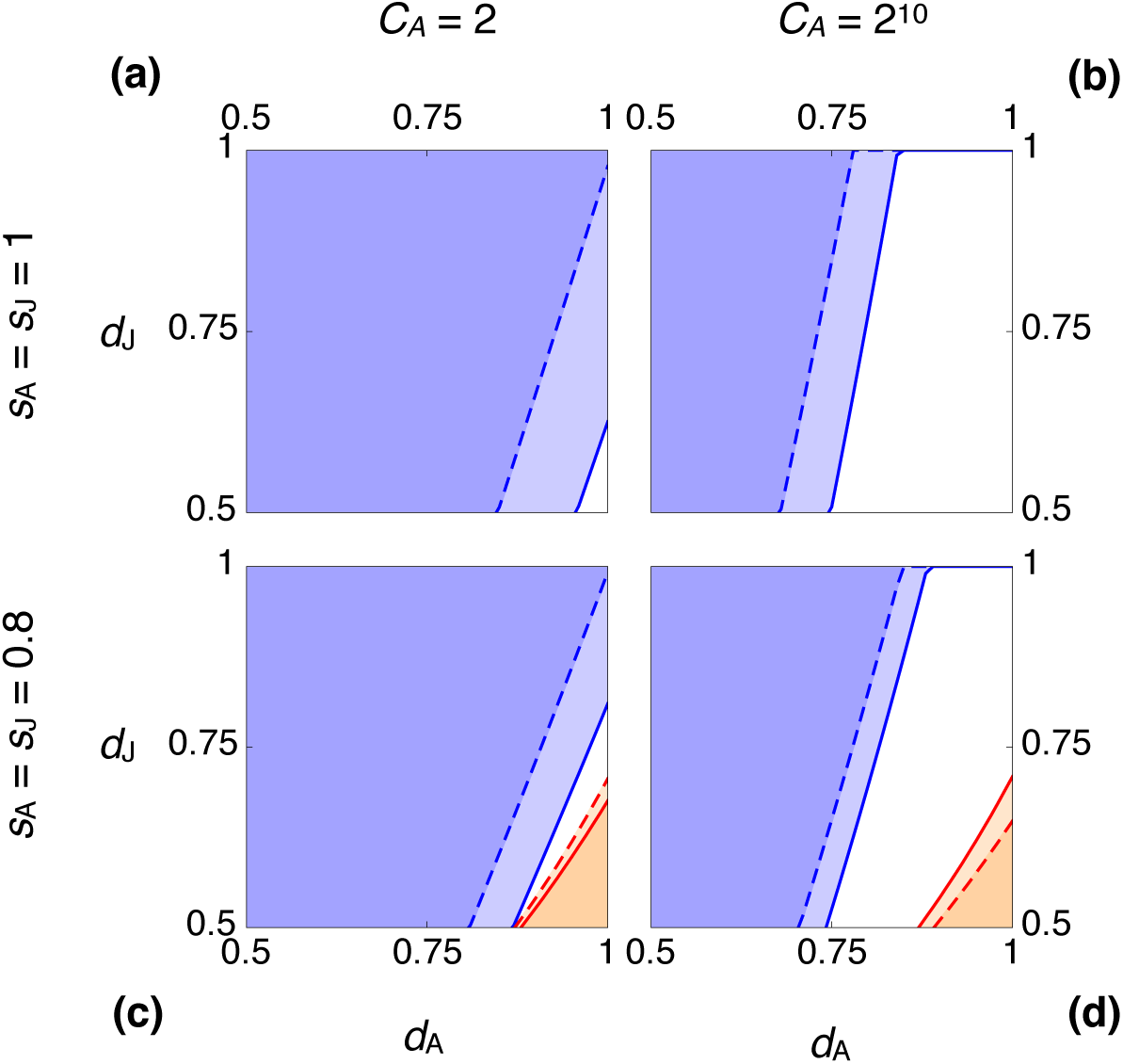
Effects of adult and juvenile dispersal survival, *d*_A_ and *d*_J_, on stability of Bequeath and Stay. Colors and curves same as in Figure 2. In each panel, horizontal axis is adult dispersal survival and vertical axis is juvenile dispersal survival. Top row: *s*_A_ = *s*_J_ = 1. Bottom row: *s*_A_ = *s*_J_ =0.8. Left column: *C*_A_ = 2. Right column: *C*_A_ = 2^10^.

### 3.5 Mixed equilibria

For the vast majority of the parameter space, either Bequeath or Stay or both are evolutionarily stable. There are no mixed, internal equilibria at which both B and S may coexist. But when adult dispersal survival is relatively high and adult baseline survival relatively low, it is possible for the orange and blue regions to overlap, for neither B nor S to be an ESS. At these parameter combinations, natural selection favors a stable mix of B and S.

Figure 5(a) shows the sensitivity analysis for adult baseline and dispersal survival, *s*_A_ and *d*_A_, fixing juvenile survival to *s*_J_ = 0.8 and *d*_J_ = 0.55. The orange and blue regions have the same meaning as in previous figures: orange indicates combinations of parameters for which Stay is not an ESS, and blue indicates combinations for which Bequeath is not an ESS. However, now there is a thin wedge where the orange and blue regions overlap. In this region of overlap, neither B nor S is evolutionarily stable. Notice that the region of overlap comprises combinations of relatively low adult baseline survival, *s*_A_, and relatively high adult dispersal survival, *d*_A_. Put plainly, when adults disperse well but survive poorly (equivalently, when juveniles disperse poorly but survive well), neither B nor S may be an ESS.

**FIGURE 5.**
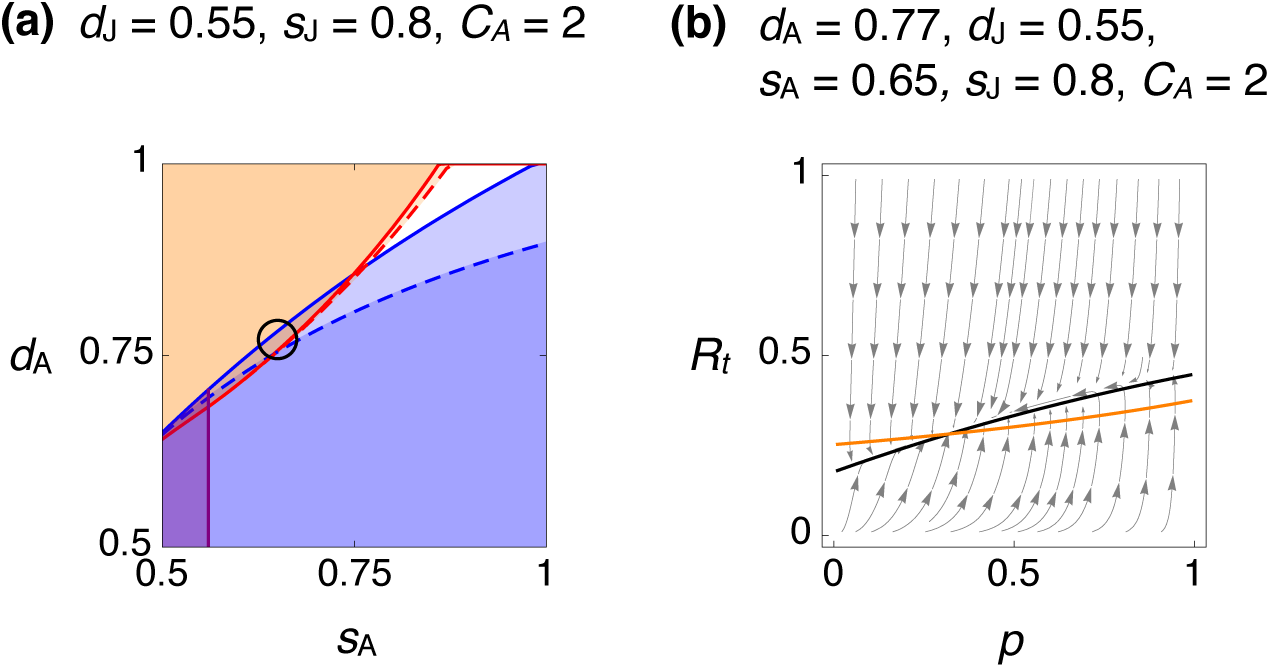
When neither B nor S is an ESS. (a) Regions in which B (blue) and S (orange) are not evolutionarily stable, as functions of adult dispersal survival *d*_A_ and adult baseline survival *s*_A_. other parameters fixed at *d*_J_ = 0.55, *s*_J_ = 0.8, *C*_A_ = 2, *ρ* = 0.5. A narrow wedge of overlap indicates combinations at which neither B nor S is an ESS. The purple region in lower-left indicates non-viable populations of S. The black circle marksthe parameter valuse used in the other panel. (b) Vector filed. The graytrajectories show the local dynamics of *R*_*t*_ and *p*. The solid black curve is the isocline for *R*_*t*_, indicating combinations of *p, R*_*t*_ for which *R*_*t*_ does not change. The orange curve is the isocline for *p*. An equilibrium lies at the intersection of the isoclines, near *p* = 0.3.

The analytical model predicts a stable mixture of B and S under these conditions. Figure 5(b) uses the fitness expressions to plot the joint dynamics of *R*_*t*_ and *p*, at the parameter values indicated by the circle inside Figure 5(a). The orange curve is the *p* isocline, combinations of *p* and *R*_*t*_ at which △*p* = 0. Below the orange curve, *p* increases. Above it, *p* decreases. The black curve is similarly the *R*_*t*_ isocline, where △*R* = *R*_*t*+1_ − *R*_*t*_ = 0. An internal equilibrium lies at the intersection of these two isoclines, near *p* = 0.3, *R*_*t*_ = 0.28.

We are not sure what to predict, given the existence of these mixed equilibria. On the one hand, this dynamic may be an unlikely outcome in natural populations, as the parameter combinations that make it possible are rare. On the other hand, nature does not randomly sample from parameter spaces. Instead, dispersal costs and baseline mortality evolve. In addition, drift may be a substantial force in natural populations, and drift will interact with selection in these models, because selection alters the habitat saturation and may decrease effective population size.

Regardless, the existence of these mixed equilibria sheds light on the general conditions that favor both B and S, and therefore aids in understanding dispersal strategy more generally. Specifically, we are struck by how hard we had to search to find mixed equilibria in these models. Unless dispersal and mortality are tuned in precise ways, selection will not favor a mix of B and S.

## 4. Discussion

We have developed and analyzed two very simple models of bequeathal. In the first, dispersal is global and random. In the second, dispersal is local and random. In both models, a single adult breeder occupies a site and produces a single juvenile offspring. Genes in the adult determine whether it evicts the juvenile, forcing it to disperse, or rather bequeaths the site to the juvenile, dispersing itself. Both adults and juveniles must compete with non-related individuals to retain or acquire breeding sites, and adults are advantaged in such competition. Adults and juveniles experience mortality during dispersal and at the end of each breeding season. Depending upon survival parameters, the habitat may or may not be saturated, but it is always uniform and static, with respect to the number and productivity of breeding sites.

Based on our results, bequeathal is most likely to be adaptive under the following conditions.

1. *In unsaturated habitat.* An unsaturated environment, with vacant breeding sites, reduces the competition a bequeathing adult faces.
2. *When adults easily defeat juveniles in contests for breeding sites.* Our models make no distinction between experience-related and size-related competitive advantages.
3. *When adults are superior to juveniles in dispersal survival.* Our models do not address whether superior survival is due to greater knowledge of the habitat or greater experience avoiding predation or even greater body size.
4. *When adults have less residual reproductive value than their offspring.* This can be true for example when an adult is less likely to survive to breed a second time than a juvenile is to survive to adulthood.

These conditions do not seem too restrictive, and indeed all of them have been suggested in the empirical literature as conditions that may favor bequeathal. As described in the *Model Results* section, these conditions are interactive and can sometimes counteract one another. Our analysis also finds many situations in which bequeathal does not evolve, even when these conditions are satisfied (for empirical examples that fail to detect bequeathal see Lambin (1997), Selonen and Wistbacka (2017)). The major reason is that bequeathal is a cooperative behavior that may impose substantial fitness costs on the adult. As a result, often even when bequeathal is adaptive—can be maintained by natural selection—it may not be able to invade the population. For most of the parameter space in our models, bequeathal is most challenged when it is rare. This positive frequency dependence creates large regions in which both bequeathal and juvenile dispersal are evolutionarily stable, making it hard to know what to predict.

Prediction is made more challenging once we remember that models of this sort are rarely valuable for their direct quantitative predictions. As the first formal models of bequeathal, these had to be simple to be productive. Despite their simplicity, they exhibit complex dynamics that demonstrate the basic tradeoffs inherent in bequeathal, tradeoffs that are likely to operate in more-complex models as well as in real populations.

### 4.1 Facultative response

The strategies we have modeled so far are inflexible. Bequeathal in nature, like other modes of resource inheritance, is more likely part of a portfolio of dispersal strategies that individuals deploy facultatively, as conditions change (Myles 1988). Models without explicit plasticity can sometimes be usefully interpreted as guides to plastic response. There are also risks that plasticity will generate novel feedback. In that case, attempting to interpret evolutionary dynamics as behavioral dynamics may frustrate and confuse. Still, it is useful to consider facultative interpretations of our results, as it helps to integrate our models with the existing literature, as well as guide future theorizing.

We have assumed that adult competitive ability, *C*_A_, is constant across individuals. If instead adults vary in competitive ability, and have some knowledge of it, then dispersal strategy may be contingent. We found that bequeathal is favored and easier to maintain when *C*_A_ is large, suggesting that larger and more aggressive individuals might do better pursuing bequeathal. There is also the possibility that individuals who already occupy a site have a prior residency advantage over immigrant intruders (Maynard Smith and Parker 1976, Kokko et al. 2006). This could apply to both both non-bequeathing adults as well as juveniles who inherit breeding sites. If such an advantage were only to apply to adults, then the conditions favoring bequeathal would be reduced.

An animal using bequeathal facultatively should be more likely to bequeath in unsaturated habitat than in a saturated one (Harris and Murie 1984, Price and Boutin 1993, Boutin et al. 1993). Unsaturated habitat favors bequeathal, because it reduces competition at an away site. Thinking ecologically, stochastic disturbance that creates new unoccupied habitat, or rather removes a large portion of the population, may encourage bequeathal. Provided adults enjoy higher dispersal survival than do juveniles, facultative bequeathal following disturbance or an increase in baseline mortality may allow a population to rescue itself. This is because habitat saturation would be higher under bequeathal than under juvenile dispersal. Such a mechanism can work in our models. If it can also function in natural populations, even rare bequeathal following disturbance may be ecologically important, because it will allow populations to persist in otherwise challenging habitats.

Bequeathal may also be a facultative strategy at end of life (Descamps et al. 2007). We found that when adults experience higher baseline mortality than do juveniles, selection tends to favor bequeathal. This is because an adult with low survival expectation has low residual reproductive value. In more complex life histories, where for example the survival probability changes with age, it might be possible that young adults will be selected to evict offspring, while older adults are selected to bequeath.

Another aspect of life history that may lead to facultative bequeathal is timing of birth (Price 1992). When females give birth late in the season, juveniles may not have sufficient time to grow to a size that would allow them to successfully disperse and compete for a breeding site. In contrast, an offspring born early in the season may have an advantage, competing against an average juvenile. If so, bequeathal may be favored late in the breeding season, even when it cannot be favored early in the season. Evidence consistent with this has been found in plateau pikas (Zhang et al. 2017).

Finally, we have treated habitat saturation as a uniform factor. In reality, local saturation matters more than global saturation. Adults who know their range and are aware of open sites may do better bequeathing, even though the same individuals might do better to evict offspring, were the local environment more saturated. Along similar lines, the models could be expanded to include a flexible search strategy during dispersal (McCarthy 1999), such that dispersers are more likely to colonize empty sites and avoid those that are occupied.

### 4.2 Future directions

Conspicuously absent strategies in our models are site sharing and floating. In the wider literature, e.g. Brown and Brown (1984), and in other models of breeding dispersal, such as Kokko and Lundberg (2001), adults may share sites with offspring. While sharing a site, offspring either postpone reproduction or reproduce at a reduced rate, while adults suffer some cost of sharing. A sharing strategy could be introduced into our models. Instead of bequeathing or evicting the offspring, the adult could allow the juvenile to remain at the natal site, a strategy seen in red squirrels (Berteaux and Boutin 2000) as well as bushy-tailed woodrats (Moses and Millar 1994). Parameters would be needed to specify juvenile and adult reproductive rates at a shared site, and unless juvenile reproductive rate is zero, some additional aspects of dispersal strategy would be needed to address conflict between offspring of both residents. In this way, the models could begin to integrate with the reproductive skew and reproductive queuing literatures (Koenig et al. 1992, Keller and Reeve 1994, Clutton-Brock 1998, Kokko and Johnstone 1999, Johnstone 2000, Cant and English 2006).

Similarly, our models could be expanded to include the possibility of floating, or waiting in interstitial habitat for breeding sites to become available (Penteriani et al. 2011). Allowing floaters would increase the average number of competitors at each site, but since this effect would be experienced at both Stay and Bequeath sites, it is unclear exactly how this would influence bequeathal, and would depend on the assumptions made about the survival and competitive abilities of floaters. Additional modeling would be needed to clarify this question.

Our models deliberately studied reproduction of a single offspring, so that we could study bequeathal in the absence of sibling rivalry and the greatly enlarged strategy space that must arise once families can be of any size. Some of the species for which bequeathal has been observed do tend to have small litters frequently with only a single offspring surviving each season (e.g., woodrats McEachern et al. (2009)). However, many animals have larger litters/broods. It may be that bequeathal is likely to be rare in species with large litters, because of reduced offspring viability, the conflicts of interest that arise among siblings, as well as an expected increase in habitat saturation. To explore these ideas, we envision an expanded strategy space in which adults both evict a certain number of offspring (from zero to all) as well as determine whether the adult itself disperses (bequeaths). The bequeathal strategy studied in this case would correspond to adult dispersal and eviction of all but one offspring from the natal site. However many other dispersal patterns would be possible within this strategy space, including total eviction with adult residency and all-but-one eviction with adult residency.

A feature of bequeathal in many species is that a durable resource—often a den, burrow, or cache—is bequeathed together with the territory. Our models ignored the construction and persistence of such resources. Presumably there is some cost of building a den, and if adults are better able to afford these costs, then our models may underestimate bequeathal’s adaptiveness. As a first sketch of a model with dynamic site resources, suppose that each site is also characterized by the presence or absence of a den. When a site has a resident, a den can be maintained. In the absence of a resident, a den has a probability of decaying. A den can be constructed at a site at a fitness cost *k*_A_ for adults and *k*_J_ for juveniles, where *k*_A_ *< k*_J_. We think this model could be analytically specified under global dispersal, generating a three dimensional dynamical system in which the frequency of bequeathal, the residency rate, and the proportion of sites with dens would all evolve together.

Our models have ignored males, treating them as ambient and causally inert. Provided that males are carriers of the bequeathal allele, and that there is no shortage of males, this assumption may be harmless. However, suppose instead that males also depend upon the same sites for survival. Then different dispersal strategies may be favored, depending upon both an individual’s sex and the sex of its offspring. As observed instances of bequeathal appear to be sex-biased towards both females (Fisher et al. 2017) and males (Banks et al. 2002), a theoretical framework that explains the conditions under which sex-biased bequeathal might evolve would be of interest and would further unite the dispersal and reproductive skew literatures. It is worth noting that there are currently no empirical examples of adult males bequeathing territory to offspring. In many mammals, this makes sense, given that males often do not co-reside with offspring or provide any parental care. Paternity uncertainty and the effect it has on relatedness may also discourage males from bequeathing territory.

Lastly, observed instances of bequeathal are heavily biased towards mammals, particularly rodents. However, there is no reason to believe that this is a uniquely mammalian phenomenon. Instances of bequeathal may be over-looked and attributed to adult mortality if the juvenile remains in the natal territory and adult movement is not tracked or detected. Bequeathal may be observed in other solitary breeding species with overlapping generations who depend on discreet resources such as dens, burrows, nests, or caches to survive– particularly if these resources are limited or costly to build. We encourage researchers studying dispersal in other taxa that fit these criteria to entertain bequeathal as an alternative dispersal hypothesis– especially in commonly known breeding dispersers like birds. This requires researches to track the relatedness of juveniles and adults in a population, location of adults and juveniles after breeding, and availability of potential territories in space across several breeding season to adequately identify and test the predictions of our model.

## Data Accessibility

A Mathematica notebook (Wolfram Research Inc. 2010) of these models, as well as R code (R Core Team 2018) for numerical simulations, can be found at this repository: https://github.com/bjbarrett/bequeathal

## ACKNOWLEDGMENTS

This paper is dedicated to the memory of Parry M.R. Clarke, our dear friend and colleague who did most of the theoretical work presented here. We thank members of the UC Davis Animal Behavior and Human Behavioral Ecology programs for their attention and advice on early versions of this work. We also thank two anonymous reviewers for their thoughtful and constructive feedback.

## AUTHOR CONTRIBUTIONS

**Parry M.R. Clarke** contributed to the theoretical analysis and interpretation, and helped draft earlier versions of the manuscript.

**Mary Brooke McElreath** contributed to the original design and concept of the study, analysis and interpretation of the models, and helped draft the manuscript.

**Brendan J Barrett** contributed to the interpretation of the models, maintenance of R code and supplemental materials, as well as the writing and revision of the final manuscript.

**Karen E Mabry** contributed to the original design and concept of the study and helped draft the manuscript.

**Richard McElreath** contributed to the original design and concept of the study, supervised the theoretical analysis and interpretation, and helped draft the manuscript.

## APPENDIX

### Inclusive fitness in the local dispersal model

Limiting our definitions to *p ∈ {*0, 1*}*:

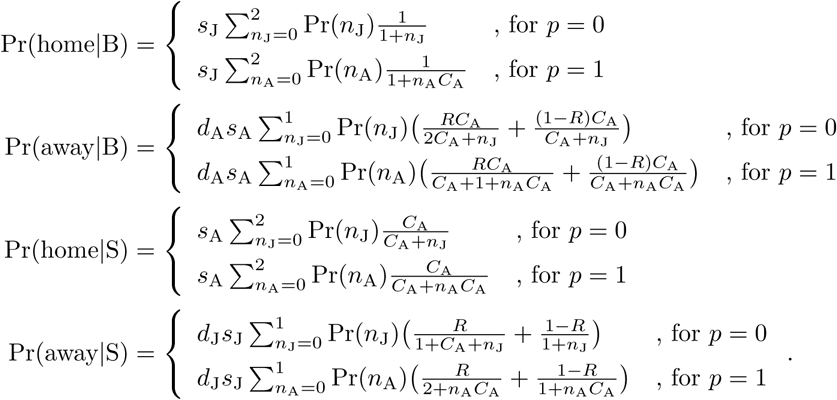

Inclusive fitness is defined identically to the global dispersal model, using the probabilities above: *W* (B) = *ρ* Pr(home*|*B) + Pr(away*|*B), *W* (S) = Pr(home*|*S) + *ρ* Pr(away*|*S).

### Bequeathal is an ESS under asexual reproduction

Let *d*_A_ = *d*_J_ = *s*_A_ = *s*_J_ = *ρ* = 1. Under these conditions, the environment will remain saturated, and so *R* = 1. Under these conditions, the average number of immigrants to each site is 1, and all dispersers are adults. Since the environment remains saturated, the average per-site success of a common strategy must be 1 (the carrying capacity). Therefore we only need to compute the S invader fitness and compare it to 1 to prove whether B is an ESS.

#### Global dispersal

The probability distribution of adults arriving to a site simplifies to a straight Poisson probability:

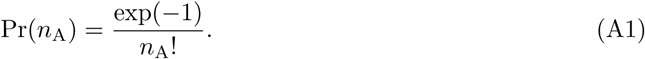

The probability that a mutant S adult retains a home site is now:

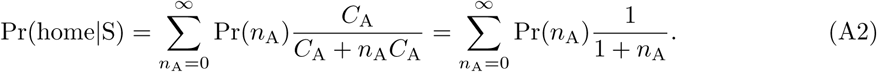

And the probability the dispersing juvenile S acquires the away site is:

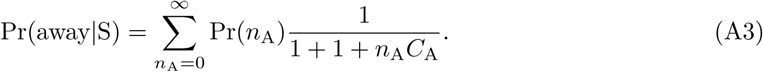

These expressions, and their sum, are not so easy to evaluate for any *C*_A_ *>* 1. But we can inspect the limits and still deduce that B is an ESS for any *C*_A_ *>* 1. First, consider when *C*_A_ = 1. Then:

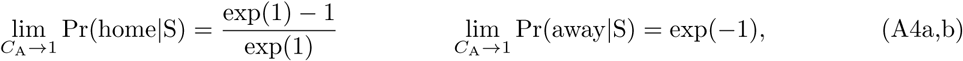

which sum to 1. So when *C*_A_ = 1, there are of course no differences between B and S strategies, so they have the same fitness. Second, consider when *C*_A_ *→ ∞*. Then:

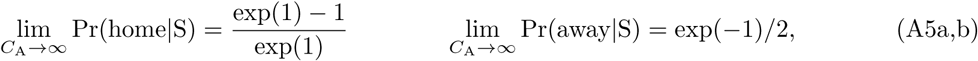

which sums to less than 1. Since the effect of increasing *C*_A_ on Pr(away*|*S) is to reduce it, B is anESS for any *C*_A_ *>* 1.

A similar argument proves that B can always invade a population of S, under the same conditions.

#### Local dispersal

The probability that a mutant S adult retains a home site is:

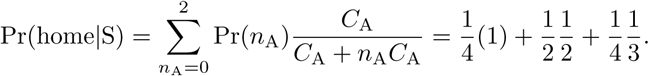

And the probability the S juvenile acquires an away site is:

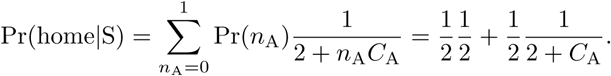

Under asexual reproduction, mutant fitness is just the sum of these two expressions. This sum is never greater than 1—resident fitness—provided *C*_A_ *>* 1. Therefore B is an ESS.

A similar argument shows that B can always invade S, under the same conditions.

### Bequeathal is an ESS under sexual reproduction, provided *C*_A_ is large enough

#### Global dispersal

When *ρ <* 1, B is not an ESS for any *C*_A_ *>* 1. But B is an ESS for *C*_A_ *→ ∞*. To demonstrate this result, assume again *d*_A_ = *d*_J_ = *s*_A_ = *s*_J_ = 1. As a result, again *R* = 1. However, now assume 0 *< ρ <* 1. Resident fitness will not be 1 now, but instead some fraction of 1, as offspring fitness is discounted by *ρ*. So we must calculate both resident and invader fitness.

A resident B juvenile retains home site with probability:

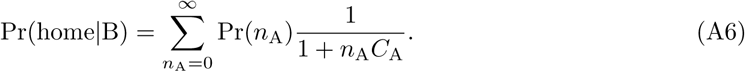

And a resident B adult acquires an away site with probability:

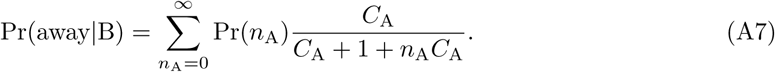

And resident B inclusive fitness is given by *W* (B) = *ρ* Pr(home*|*B) + Pr(away*|*B). Invader fitness is as in the previous section, but with inclusive fitness *W* (S) = Pr(home*|*S) + *ρ* Pr(away*|*S). Consider first when *C*_A_ = 1. Taking limits, resident fitness is:

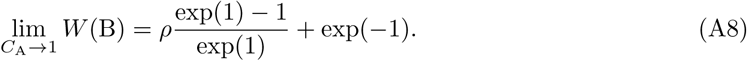

And likewise for the invader:

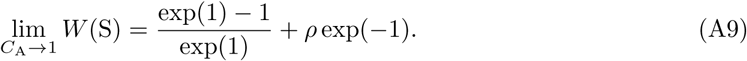

Since (exp(1) − 1)*/* exp(1) *>* exp(*-*1), *W* (S) *> W* (B) for any *ρ <* 1. Therefore B is never an ESS, when *C*_A_ = 1 and *ρ <* 1. A similar argument demonstrates that S is always an ESS under the same conditions.

Now consider when *C*_A_ *→ ∞*. Again, taking limits:

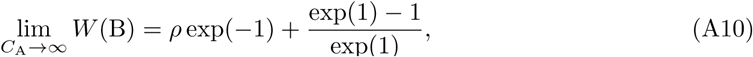

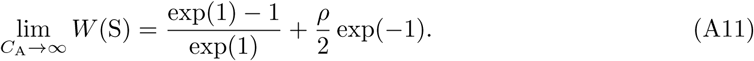

And now *W* (B) *> W* (S) for any *ρ >* 0.

Therefore B is an ESS, once *C*_A_ is sufficiently large. We cannot prove analytically how large *C*_A_ must be to cross the threshold required to make B an ESS. But we can be sure such a threshold exists, as the effect of *C*_A_ on the probabilities of winning sites is monotonic.

#### Local dispersal

In the case of local dispersal, an exact condition can be derived. B is an ESS, provided:

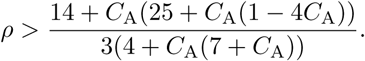

Unfortunately, nothing can be gained by inspecting this inequality directly, aside from noting that greater relatedness favors Bequeathal. This inequality defines the dashed blue boundary in Figure 2(a).

